# Selective breeding enhances coral heat tolerance even over small spatial scales

**DOI:** 10.1101/2025.06.04.657771

**Authors:** Alexandra Kler Lago, Kate Kiefer, Marie E. Strader, Teresa Baptista Nobre, Stephanie F. Hendricks, Claudio Richter, Christian Wild, Kate Quigley

## Abstract

Coral reefs globally are experiencing escalating mass bleaching and mortality. Reefs along the western Indian Ocean have been relatively unimpacted. We assessed heat tolerance baselines in two widespread reef-building *Acropora* species *a*nd used selective breeding from two thermally distinct (present day and stress histories) northern (Mean Monthly Maximum 27.9 °C) and southern (26.6 °C) reefs along the Ningaloo World Heritage Reef. Fitness responses were measured in control and heat stress temperatures (adults = 31.0 ^°^C, larvae = 35.5 ^°^C), including survival, tissue necrosis, bleaching, and photosynthesis. Larvae with one parent from the warmer population exhibited >2.2-fold higher survival under heat stress, while those with both parents from the warmer population survived 1.6-fold better (compared to control larvae with two parents from the cooler population). Photosynthesis was maintained in both species and both populations, suggesting heat responses were host driven. Adults from both populations from one species (*A. tenuis*) exhibited similar responses to heat, while the other (*A. millepora*) was more variable. These findings are the first to demonstrate that selective breeding can provide heat tolerance enhancement for corals in the Indian Ocean and will be critical to preparing for future marine heatwaves.

## Background

Since the preindustrial era (∼1850-1900), coral reefs have experienced four global mass bleaching events (1). This has caused the global decline of approximately 50% in coral cover, mainly attributed to an on average +1 ^°^C of ocean warming and intensified marine heatwaves (MHWs) (2). Within the next three decades, climate predictions strongly indicate an increase in ocean surface temperatures by +1.5 to +2.0 ^°^C, accompanied by warmer (+1.9 to +2.5 ^°^C), prolonged, and more frequent (4.1x to 5.6x) MHWs per decade (3, 4). This warming will likely result in annual mass bleaching across most reefs, with a projected 70% to 99% loss of coral cover and potential ecological collapse beyond +2.5 ^°^C warming (3-4). Such extensive mortality indicates that the rate of temperature increase may be outpacing the natural rate of thermal adaptation in corals, which will impact coral populations recovery and replenishment (5,6). Given coral reefs’ ecological and economic importance (7), active interventions beyond conventional reef management and restoration approaches are urgently needed. These actions should bolster resilience to maintain critical ecosystem functions and services until greenhouse gas emissions and warming are brought under control (8).

Emerging active intervention tools like assisted evolution have been proposed to accelerate adaptation by increasing heat tolerance in corals and their symbionts faster than natural rates (8). These tools include selective breeding of the coral host (9), which consists of selecting and reproductively crossing coral parental stocks with heritable fitness-related traits associated with higher heat tolerance to enhance these same traits in the offspring. While there are multiple approaches to selecting parental stocks, which vary with scalability, costs and technical dependency, most studies have used local summer maxima temperatures and/or bleaching responses as proxies for heat tolerance in parental stocks for breeding (10). Another study also found specific historical temperature, daily temperature variation and thermal stress metrics for predicting selection of thermally tolerant parental stocks (11). The end goal is to increase desired traits (e.g., increased heat tolerance) whilst maintaining the genetics of local populations (12). When combined with movement, transferring selected offspring to at-risk reef locations — a process known as Assisted Gene Flow — can improve the resilience of coral populations within their known ranges (12).

Selective breeding studies, which have thus far primarily been on the Great Barrier Reef (GBR) and the Persian Gulf, have achieved promising results in selected offspring (i.e., larvae and juveniles) with an increased survivorship to warming (13-15). This adaptive response is largely due to moderate to high heritability of heat tolerance across multiple life stages and symbiont associations (e.g., narrow-sense heritability h^2^ > 0.5: 33-35). Importantly, heritability across many traits in corals’ is considerably heterogeneous across stages/ages, growth forms, and environments (16). This variation underscores the importance of generating baseline information on heat tolerance and heritability across diverse coral populations for the development of selective breeding strategies.

In the last three decades, the Ningaloo World Heritage area in Western Australia (WA), one of the world’s most remote, diverse and extensive fringing reef systems spanning ∼290 km, has experienced five recorded severe heat stress events (Degree Heating Weeks (DHW) > 8 ^°^C-weeks) since 1998 (17). This includes the severe 2011 MHW (18) which resulted in sea surface temperatures (SST) up to 5 ^°^C warmer than average for more than 10 weeks (17, 19, 20). Ningaloo harbors half of the coral species in the Indian Ocean (21), making it an outstanding ark of global coral biodiversity and a target for conservation priority under the risk of annual bleaching by 2050 (22, 23). During these past heat stress events, coral bleaching and mortality responses have been spatially variable with some reports showing an up to 92% and 32% decline in the northeastern and southern areas of Ningaloo, respectively, while coral cover remained stable in the northern region (19, 24, 25). Thus far, baselines in coral heat tolerance have not yet been determined and it is unclear how vulnerable these populations are to climate change. Further, no selective breeding studies have been conducted in this globally important region.

To address these knowledge gaps, we conducted assessments of breeding feasibility and heat tolerance for two common and widespread Acropora species sourced from the northern and southern regions of Ningaloo Coast. These two locations on the reef were selected for their distinct thermal profiles (Figures 1 and 2). We tested and quantified multiple fitness-related responses (i.e., survival, tissue necrosis, bleaching, and photochemical efficiency) in adult (parental) corals and their selectively bred larval offspring exposed to experimental heat stress. We assessed survival under heat stress of larval offspring produced from northern and southern corals as well as from a mix of these populations. Overall, our study demonstrated that selective breeding significantly improved the heat tolerance of an early life-history stage in the two Acropora coral species by up to 2.2-fold, even though parental responses to heat stress were variable.

**Figure 1.**
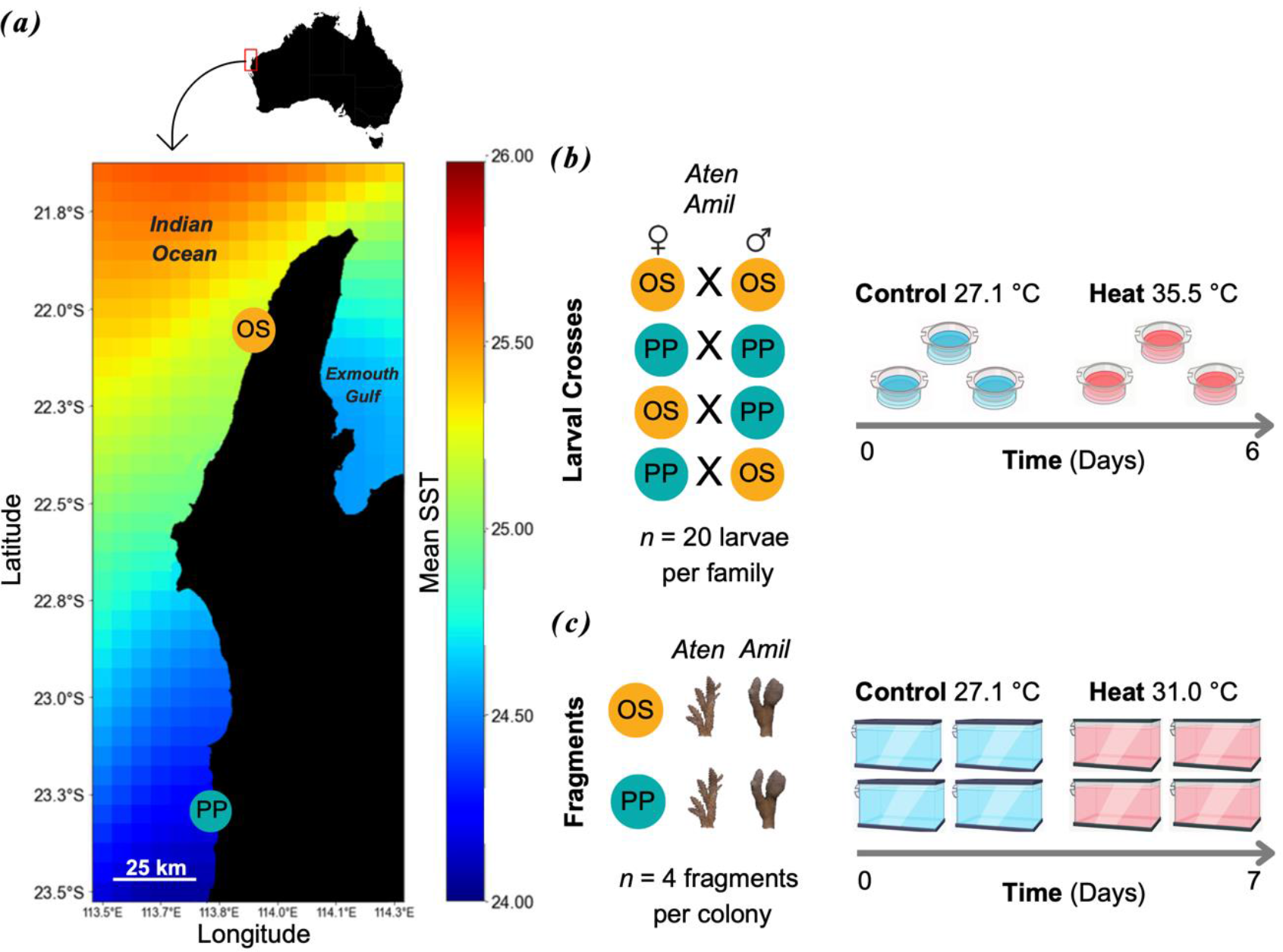
Experimental design for adult and larval heat stress. (a) Mean annual sea surface temperatures (SST) along the Ningaloo Coast (1985–2022; NOAA CoralTemp). Collection sites of *Acropora tenuis* (Aten) and *Acropora millepora* (Amil) at Oyster Stacks (OS, yellow) and Pelican Point (PP, blue). (b) Larval heat stress: 20 larvae per family from intrapopulation (OS×OS, PP×PP) and interpopulation (OS×PP, PP×OS) crosses were tested at 27.1°C and 35.5°C. (c) Adult heat stress: 3-4 fragments per genotype from both populations were tested at 27.1°C and 31.0°C with daily assessments of survival, necrosis, bleaching, and effective quantum yield.

**Figure 2.**
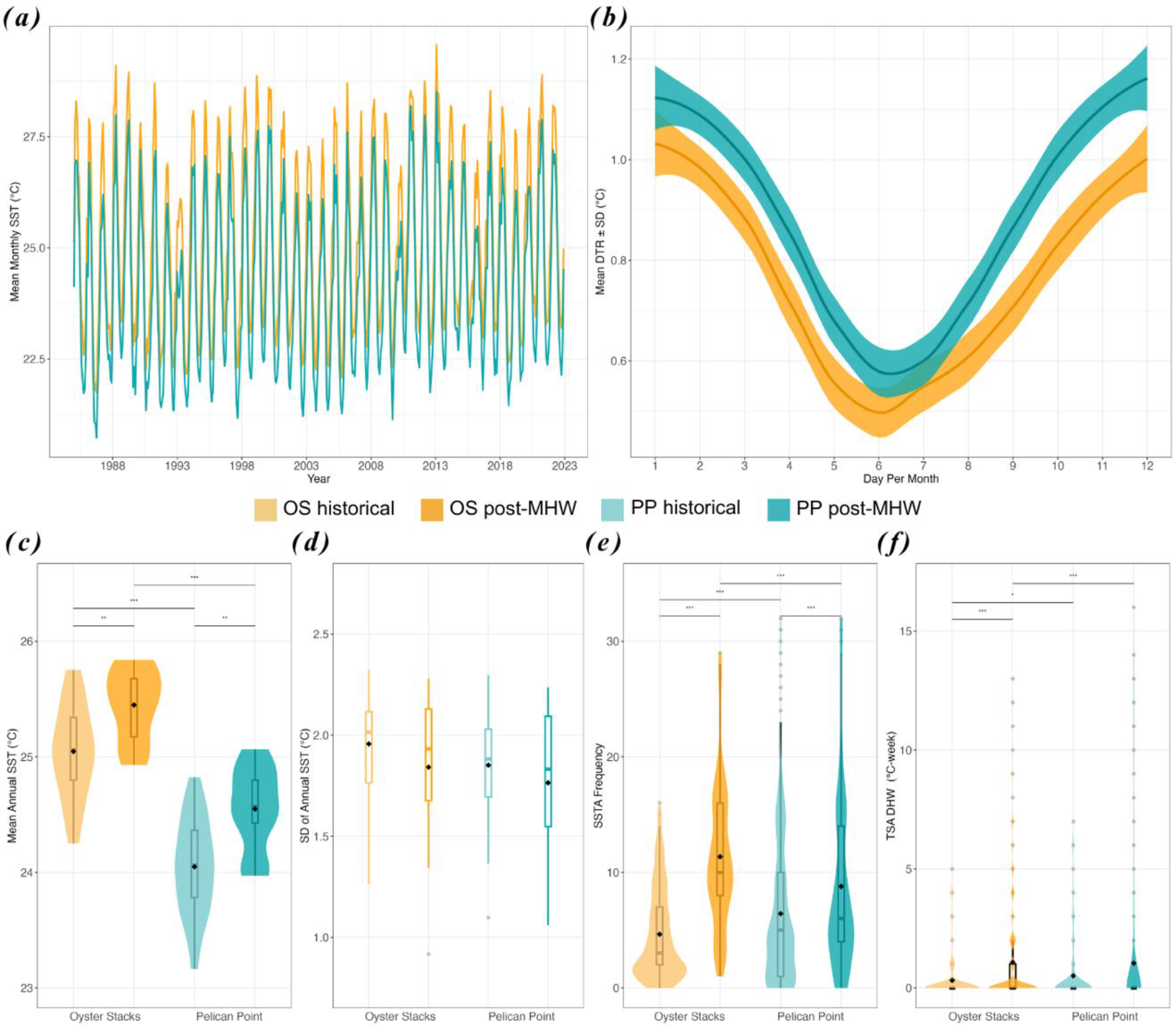
Temperature metrics of collection sites on Ningaloo. (a) Mean monthly SST time series (1985–2023) for Oyster Stacks (OS, yellow) and Pelican Point (PP, blue). (b) Diurnal temperature range ± standard deviation (DTR ± SD) over a 12-month cycle (2016–2022). (c) Mean annual SST (SST_mean) for historical (1985–2010, pale) and post-MHW (2010–2022, dark) periods. Panels (d–f) show annual SST standard deviation (SST_stdev), sea surface temperature anomaly frequency (SSTA_Freq), and cumulative heat stress in degree heating weeks (TSA_DHW). Boxplots display medians (center lines), quartiles (box limits), 1.5× interquartile range (whiskers), and outliers (points); diamonds indicate means. Horizontal lines with asterisks represent significant differences (* *P* < 0.05, ** *P* < 0.01, *** *P* < 0.001) between locations and periods (Wilcoxon’s test). *P* values are detailed in SI Table 5.

## Results

### Distinct thermal profiles between reefs in historical and heat stress periods

Temperature metrics related to SST (SST_mean, SST_stdev) and thermal anomaly (SSTA_Freq, TSA_DHW) were assessed for OS and PP during two periods: historical (1985-2010) and post-MHW (2010-2022), which includes the heat stress events from 2011 to 2013.

During the historical period, both locations had similar annual temperature cycles, with April being the warmest month (Maximum Monthly Mean: MMM) (M ± SD = 27.8 ± 0.9 ^°^C for OS; 26.5 ± 1.0 ^°^C for PP), and September the coolest. OS recorded a significantly higher SST_mean than PP by +1 °C (OS: 25.0 ± 0.4 °C, PP: 24.0 ± 0.4 °C) (Figure 2c, Wilcoxon, *P* < 0.001). SST_stdev was similar between sites (OS: 2.0 ± 0.3, PP: 1.9 ± 0.3) (Figure 2d, Table S5, Wilcoxon, *P* = 0.174). Thermal anomalies showed OS had a lower SSTA_Freq (OS: 4.6 ± 3.9, PP: 5.3 ± 5.1) and TSA_DHW (OS: 0.3 ± 1.0, PP: 0.6 ± 1.4) (Figure 2e, f, Table S5, Wilcoxon, *P* < 0.001, *P* = 0.018 for SSTA_Freq and TSA_DHW, respectively).

In the post-MHW period, both sites exhibited a significant rise in SST_mean compared to historical baselines: OS increased to 25.4 ± 0.3 °C (SI Table 5, Wilcoxon, *P* = 0.004), and PP to 24.5 ± 0.4 °C (Table S5, Wilcoxon, *P* = 0.002). SST_mean remained significantly different between the two sites (Table S5, Wilcoxon, *P* < 0.001). MMM also increased in a similar way (28.1 ± 0.7 ^°^C for OS; 26.9 ± 0.8 ^°^C for PP). SST_stdev remained stable (Table S5, Wilcoxon, *P* = 0.527, *P* = 0.586 for OS and PP, respectively) with non-significant decreases (OS: 1.8 ± 0.4 ^°^C, PP: 1.8 ± 0.4 ^°^C, Table S5, Wilcoxon, *P* = 0.511).

Despite similar variation in mean annual temperatures and close relative geographical proximity, OS and PP were distinct in their high-frequency temperature variability. PP recorded a significantly higher mean DTR than OS (OS: 0.8 ± 0.5 ^°^C, PP: 0.9 ± 0.5 ^°^C) (Figure 2b, Table S5, Wilcoxon, *P* < 0.001). Monthly DTR variations were significant between locations (Table S5, Wilcoxon, *P* < 0.001), except for January and February (*P* = 0.246, *P* = 0.154, respectively). Maximum DTR occurred in December (OS: 1.0 ± 0.7 ^°^C, PP: 1.2 ± 0.5 ^°^C) and minimum in June (OS: 0.5 ± 0.4 ^°^C, PP: 0.6 ± 0.2 ^°^C).

Post-MHW, both locations saw a significant increase in thermal anomalies. OS had 2.5× higher SSTA_Freq (11.4 ± 6.1) and PP 1.7× (8.8 ± 7.5) (Figure 2e, Table S5, Wilcoxon, *P* < 0.001). OS also had significantly more anomalies per year (2.7 weeks) than PP (Wilcoxon, Table S5, *P* < 0.001). TSA_DHW increased for both sites; OS from 0.7 DHW to 1.1 DHWs ± 2.3 (Table S5, Wilcoxon, *P* < 0.001) and PP from 0.5 DHW to 1.0 DHWs ± 2.9 (Table S5, Wilcoxon, *P* = 0.238). OS had significantly higher mean TSA_DHW relative to PP (Table S5, Wilcoxon, *P* < 0.001), indicating a higher frequency and magnitude of thermal anomalies compared to PP.

### Differences in selected larval survival responses to heat stress between reproductive crosses

*A. tenuis* showed a significant difference in larval survival for most families produced (SI Table 6 for *t*-test *P* values) and for all reproductive crosses between temperature treatments after 6 d of experimental heat stress (Figure 3a, Figure S3a-d and Table S7, Log-Rank KM, *Ps* < 0.001). In the control treatment, there was no difference in *A. tenuis* larval survival between crosses, except for one (Table S7, Log-Rank KM, *P* = 0.160 for OS×OS vs. OS×PP). However, in heat conditions, larval survival differed significantly between all crosses (SI Table 7, Log-Rank KM, *P* < 0.001 for all pairwise comparisons). In heat conditions, *A. tenuis* larvae from intrapopulation crosses recorded a mean endpoint percent survival of 71.9 ± 17.3% (OS×OS) and 44.7 ± 19.0% (PP×PP), while those from interpopulation crosses were 66.2 ± 4.1% (OS×PP) and 52.6 ± 4.0% (PP×OS), which were 1.4× to 2.2× lower compared to controls (OS×OS: 98.6 ± 3.9%, PP×PP: 96.1 ± 4.7%, OS×PP: 98.3 ± 1.1%, PP×OS: 96.7% ± 0.9) (Figure 3b). Relative to the intrapopulation PP×PP larvae, which recorded the lowest mean endpoint survival, intrapopulation OS×OS larvae had the highest survival gain by 1.6× (+27.2%), followed by interpopulation crosses OS×PP with 1.5× (+21.5%) and PP×OS with 1.2× (+7.9%) under heat conditions (Figure 3b). Within each cross, there was a large variation in mean endpoint survival between larval families from 31.7% in OS×PP up to 58.3% in OS×OS (Figure 3a).

**Figure 3.**
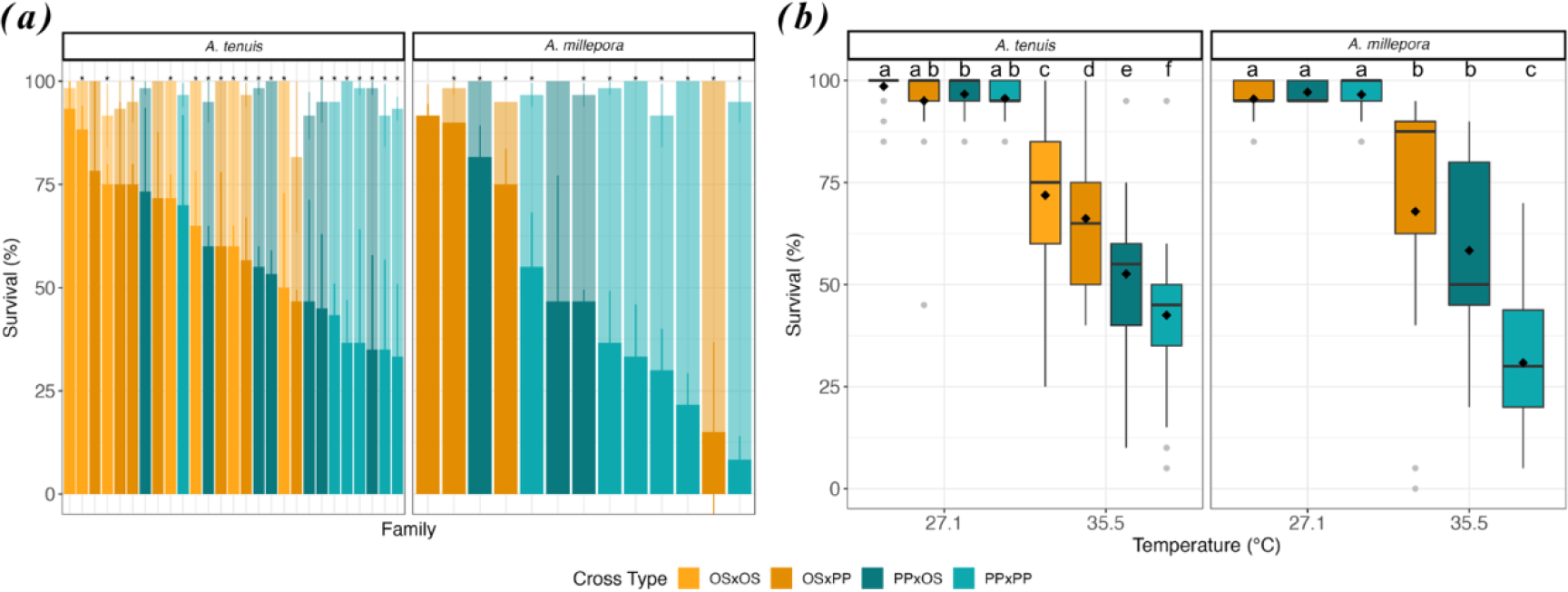
Survival responses in selected larvae under heat stress. (a) Variation in percent survival (M ± SE) among *Acropora tenuis* (*n* = 3,320) and *Acropora millepora (n* = 1,560) larval families from intrapopulation and interpopulation crosses after 142 and 93 hours of exposure to 35.5°C (foreground) versus 27.1°C (background). (b) Median survival in larval crosses exposed to control (27.1°C) and heat (35.5°C) treatments. Boxplots show medians (center lines), quartiles (box limits), 1.5× interquartile range (whiskers), and outliers (points); diamond shapes indicate the mean survival. Asterisks indicate significant differences between treatments using one-tailed *t*-tests with adjusted *P* values (SI Table 6). Letters denote significant differences across all treatment combinations within each species based on Kaplan-Meier Log-Rank tests with Bonferroni-corrected *P* values (SI Table 7).

For *A. millepora*, there was also a significant effect of temperature in larval survival for most families (Table S6 for *t*-test *P* values) and for all reproductive crosses after 4 d of heat exposure (Figure 3a, SI Figure 3e-g and Table S7, Log-Rank KM, *Ps* < 0.001). In the control, there was no difference in *A. millepora* larval survival between all crosses (see Table S7 for *P* values). In comparison, at heat, there were significant differences in *A. millepora* larval survival between intrapopulation and interpopulation crosses (Table S7, Log-Rank KM, *P* < 0.001 for all pairwise comparisons) but not between interpopulation crosses (Table S7, Log-Rank KM, *P* = 1.000 for OS×PP vs. PP×OS). In heat conditions, *A. millepora* intrapopulation cross PP×PP recorded a mean endpoint survival of 30.8 ± 4.1% while interpopulation crosses OS×PP and PP×OS had 67.9 ± 9.9% and 58.3 ± 7.9%, respectively, which were 1.9× to 3.2× lower compared to control (PP×PP: 97.8 ± 1.4%, OS×PP: 96.3 ±1.5%, PP×OS: 100.0 ± 2.2%) (Figure 3b). Relative to the intrapopulation PP×PP larvae with the lowest mean survival, interpopulation crosses had a survival gain by 2.2× (+37.1%) and 1.9× (+27.5%) (Figure 3b). Here, there was also a large variation in mean endpoint survival between larval families within each cross, from 35.0% in PP×OS up to 76.6% in OS×PP (Figure 3a).

### Differences in adult physiological responses to heat stress between reefs Survival

After 7 d of heat exposure for adult *A. tenuis* fragments, there was a significant effect of temperature on the survival probabilities across populations (Figures S2a, b and Table S7, Log-Rank KM, *Ps* <

0.001 for OS and PP). However, there was no population origin effect on the survival of *A. tenuis* fragments in both temperature treatments (Table S7, Log-Rank KM, *P* = 0.086, *P* = 0.613 for control and heat, respectively). In the heat treatment, OS fragments survived on average 2.1× less (47.5 ± 50.6%) and PP fragments 1.7× less (55.4 ± 50.0%) compared to fragments in the control (OS: 100.0 ± 0%, PP: 93.3 ± 25.1%) (Figure 4a).

**Figure 4.**
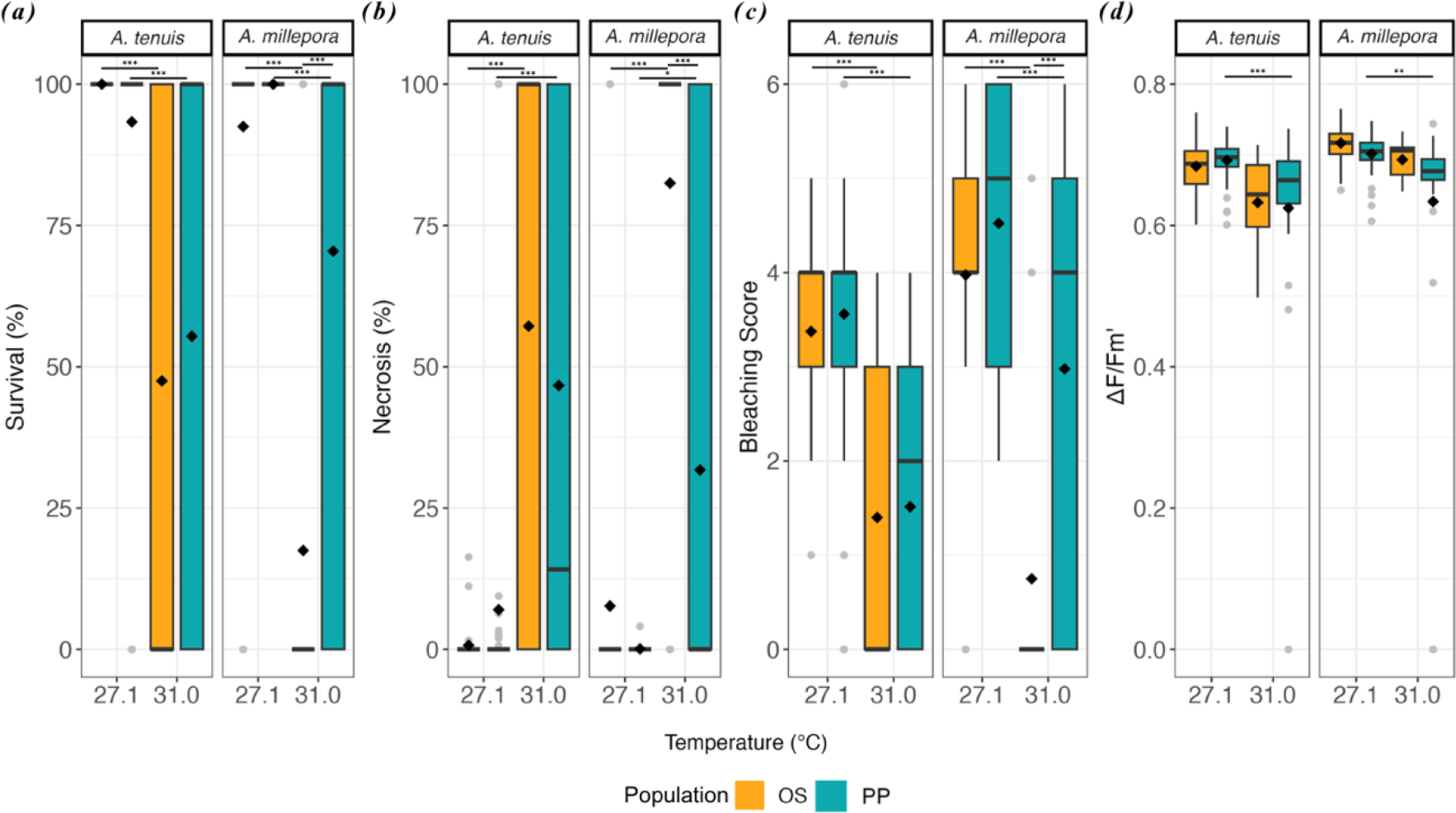
Median physiological responses of adult coral fragments under heat stress. Endpoints of (a) percent survival, (b) percent necrosis, (c) bleaching score, and (d) effective quantum yield (ΔF/Fm’) in *Acropora tenuis* (*n* = 229) and *Acropora millepora* (*n* = 168) fragments from OS and PP after 7 days of exposure to control (27.1°C) and heat (31.0°C) treatments. Boxplots show medians (center lines), quartiles (box limits), 1.5× interquartile range (whiskers), and outliers (points); diamond shapes indicate means. Horizontal lines with asterisks denote significant differences (* *P* < 0.05, ** *P* < 0.01, *** *P* < 0.001) determined by Kaplan-Meier Log-Rank tests for survival and GLMMs Tukey post hoc tests for other responses. Adjusted *P* values (Bonferroni correction) are detailed in the text.

For *A. millepora* fragments, temperature also had a significant effect on the survival probability for each reef at end of the heat stress treatment (Figures S2c, d and Table S7, Log-Rank KM, *Ps* < 0.001 for OS and PP). Compared to *A. tenuis*, population origin had a significant effect on the survival probability of *A. millepora* fragments in the heat treatment (Table S7, Log-Rank KM, *P* < 0.001). At the end of the heat exposure, OS fragments survived on average 5.7× less (17.5 ± 38.4%) and PP fragments 1.3× less (70.5 ± 46.2%) than fragments in the control (OS: 100.0 ± 0%, PP: 92.5 ± 26.7%) (Figure 4a). Between populations, PP fragments had a mean endpoint survival at heat 4.0× higher than that of OS fragments (Figure 4a).

### Tissue necrosis

For *A. tenuis* fragments, there was a significant difference in tissue necrosis owing to temperature across populations at the endpoint of heat exposure (Figure 4b and Tables S8 and S9, Tukey post-hoc beta GLMM, *Ps* < 0.001 for OS and PP). However, population origin did not have a significant effect on necrosis in both temperature treatments (Figure 4b and SI Tables 8 and 9, Tukey post-hoc beta GLMM, *Ps* =1.000 for heat and control). Relative to average tissue loss in the control (OS: 0.7 ± 3.1%, PP: 7.0 ± 25.1%), OS fragments of *A. tenuis* lost 78.4× more tissue (57.2 ± 47.3%) whereas PP fragments experienced 6.7× more tissue loss (46.7 ± 48.8%) under heat stress (Figure 4b).

Similarly for *A. millepora*, temperature had a significant effect on necrosis of fragments across populations at the endpoint of heat exposure (Figure 4b and Tables S8 and S9, Tukey post-hoc beta GLMM, *P* < 0.001, *P* = 0.035 for OS and PP, respectively). Tissue loss of fragments also differed significantly between the two populations in the heat treatment (Figure 4b and Tables S8 and S9, Tukey post-hoc beta GLMM, *P* < 0.001). At the end of the heat treatment, OS fragments of *A. millepora* recorded on average 10.7× more tissue loss (82.5 ± 38.5%) whereas PP fragments lost 353.3× more tissue (31.8 ± 45.6%) than those in control (7.7 ± 26.7% for OS, 0.1 ± 0.6% for PP) (Figure 4b). Between populations, PP fragments recorded 2.6× less tissue loss than those from OS at heat (Figure 4b).

For the fragments that died during the experiment, these fragments exhibited rapid tissue loss from 0% to 100% between 24 to 48 h of experimental heat stress, however averages and differences in time to complete tissue loss were not incorporated above (Figure S4).

### Bleaching

After *A. tenuis* fragments were exposed to heat stress, bleaching scores were significantly different due to temperature (Figure 4c and Tables S8 and S9, Tukey post-hoc nb GLMM, *P* < 0.001) but not due to population origin (Figure 4c and Tables S8 and S9, Tukey post-hoc nb GLMM, *P* = 1.000). At heat, OS fragments bleached, on average, 2.4× more (1.4 ± 1.7) and PP fragments 2.4× more (1.5 ± 1.5) than those in the control (OS: 3.4 ± 0.9, PP: 3.6 ± 1.2) (Figure 4c).

For *A. millepora*, there was a significant effect of both temperatures (Figure 4c and Tables S8 and S9, Tukey post-hoc nb GLMM, *P* < 0.001, *P* = 0.001 for OS and PP, respectively) and population origin (Figure 4c and Tables S8 and S9, Tukey post-hoc nb GLMM, *P* < 0.001) on the mean bleaching scores of fragments at the endpoint of heat stress exposure. Relative to bleaching in the control (OS: 4.0 ± 1.4, PP: 4.5 ± 1.4), OS fragments bleached, on average, 5.3× more (0.8 ± 1.7) whereas PP fragments 1.5× more (3.0 ± 2.3) in the heat treatment (Figure 4c). Between populations, PP fragments recorded 3.9× higher bleaching scores compared to OS fragments at heat (Figure 4c).

### Photochemical efficiency

For *A. tenuis*, the photochemical efficiency response, as measured by the effective quantum yield (ΔF/Fm′), only decreased significantly for PP fragments in the heat treatment compared to the control at the endpoint (Figure 4d and Tables S8 and S9, Tukey post-hoc GLMM, *P* < 0.001). Mean yields at heat were 1.1× lower relative to control conditions (Figure 4d). Reef origin was not a significant factor in explaining differences in ΔF/Fm′ across temperature treatments (Figure 4d and Tables S8 and S9, Tukey post-hoc GLMM, *Ps* = 1.000 in control and heat). At the experiment endpoint, fragments across populations recorded similar mean yields under heat (OS: 0.6 ± 0.1, PP: 0.6 ± 0.2) and control conditions (OS: 0.7 ± 0.0, PP: 0.7 ± 0.0) (Figure 4d).

Similar to *A. tenuis*, only PP fragments of *A. millepora* had a significant difference in their ΔF/Fm′ due to temperature at the endpoint of heat stress exposure (Figure 4d and Tables S8 and S9, Tukey post-hoc GLMM PQL, *P* = 0.004). Also, the population origin effect on the ΔF/Fm′ of fragments was not significant (Figure 4d and Tables S8 and S9, Tukey post-hoc GLMM PQL, *P* = 1.000). Endpoint yields were similar across populations in both temperature treatments (OS: 0.7 ± 0.0 at heat, 0.7 ± 0.0 at control, PP: 0.7 ± 0.0 at heat, 0.6 ± 0.2 at control) (Figure 4d).

## Discussion

This study presented the first evidence for enhanced heat tolerance in *Acropora* offspring using only one generation of selective breeding at Ningaloo World Heritage reef. These results confirm the feasibility of selective breeding along even a small but significant thermal gradient. Interestingly, coral adults exhibited a more complex response to heat stress, less variation between warmer and cooler populations, and distinct levels of tolerance responses between coral hosts and symbionts.

### Evidence of heat tolerance boost in *Acropora* larvae

Adaptation can occur in response to selection imposed by local environmental conditions, particularly thermal regimes, across different spatial scales (26). The resulting standing genetic variation in coral populations is critical for selectively breeding corals for fitness-related traits (9,). In this study, we reported 1.6× greater heat tolerance among *A. tenuis* intrapopulation larvae from the warmer northern reef compared to those from the cooler southern reef. Additionally, these results demonstrate the potential to significantly enhance heat tolerance over even small spatial scales and a temperature difference of ∼1 ^°^C (for both their MMM and mean annual temperatures). For comparison, this temperature differential is less than the average 2 ^°^C reported in previous selective breeding experiments (11, 48, 49). In this context, selective breeding using northern parental stock sourced from a location of significantly higher mean annual temperatures and more frequent and intense recent thermal stress (i.e., post-MHW) compared to southern corals, will likely be effective in enhancing heat tolerance in this early life-history stage in the short term.

Selective breeding has been shown to significantly increase heat tolerance in critical early life-history stages of corals by mixing gametes from coral sourced from different thermal environments (9). Consistent with these past results, we found that crossing gametes with one parent from a warmer and more impacted reef with a less warm, impacted reef yielded larvae that displayed increased tolerance by up to 1.5-fold (*A. tenuis)* or 2.2-fold (*A. millepora*) compared to control larvae with both parents sourced from the cooler, less impacted reef. Translated to DHW, these surviving, selectively bred larvae showed a boost of 2.8–5.2 DHW above the expected summer maxima by 2100 (moderate RCP4.5, +1.4–2.6°C, 6) (Table S10). These findings are consistent with previous studies across multiple species and locations showing that heat tolerance is a heritable trait (14, 13, 15, 28, 29). Furthermore, the enhanced survival observed here aligns with the broad range of survivorship gains previously documented (∼ +20-70% survival) observed in selected offspring of *Acropora, Montipora, and Platygyra* species, primarily from the GBR and the Persian Gulf (23). These results corroborate the growing body of evidence that enhanced heat tolerance is possible to achieve in early life-history stages using one cycle of selective breeding. Importantly, there is growing evidence that this success is possible even across very small spatial scales, including along less than 100km along the Ningaloo or across reefs in Palau (29). Combined with breeding, these results suggest that Assisted Gene Flow could be used as a conservation strategy along the Ningaloo even across small spatial scales.

### Species-specific parental effects on larval heat tolerance

Parental genotypes play a crucial role in shaping offspring fitness and stress responses, including heat tolerance, where both the maternal and paternal identities influences offspring fitness (14, 30, 31). In our study, we found significant species-specific parental effects on the heat tolerance of interpopulation coral offspring. For *Acropora tenuis*, maternal colonies from the warmer reef conferred greater heat tolerance to offspring than paternal colonies from the same environment, aligning with a previous study on maternal influence on survival and symbiont acquisition in *Acropora* (14, 15, 29). In contrast, *A. millepora* exhibited no difference in heat tolerance between offspring with maternal or paternal colonies from the warmer reef. Other studies have shown regional and species variations in parental effects on heat tolerance, including strong paternal effects in *Platygyra* from the Persian Gulf (14) and pronounced maternal effects in *Montipora* from Hawaii (54). Taken together, these findings suggest that parental contributions to heat tolerance are highly species-and population-specific, and are likely influenced by both environmental and genetic factors. Importantly, they highlight that conservation strategies will likely need to be tailored to specific species. This emphasizes the need for selective breeding programs that consider species-specific variability when optimizing for how to enhance coral tolerance to climate change.

### Distinct heat tolerance between adult corals and selected larvae

Following heat stress events between 2011 and 2013, coral populations in northern Ningaloo were more impacted and showed greater recovery compared to southern populations (17, 25, 33). This suggests that northern populations have already undergone some selection for increased tolerance to due to disturbance and could explain the higher tolerance transfer from northern parents to offspring, even if northern and southern adult showed similar heat tolerance. This difference between adult and offspring tolerance may be due to their symbiotic states (aposymbiotic larvae and symbiotic adults) and not necessarily surprising given it aligns with findings where adult *Acropora* GBR populations showed reduced variation in tolerance compared to their selected offspring (29). Other studies also found variable stress responses across life-history stages - with less tolerant parents yielding more tolerant offspring (e.g., 34-36). Other factors, like symbiosis, transgenerational plasticity, and parental environmental provisioning (e.g. lipids) play a significant role in determining heat tolerance aside from the host genetics (37). Differences between larvae and adults here suggest that symbionts and non-genetic effects may play an important role in determining heat tolerance in *Acropora* corals in Ningaloo and further work is needed to tease apart the relative contribution of both environmental and genetic effects on heat tolerance patterns.

### Host-driven acute heat stress responses in adult corals across reefs

The underlying heat tolerance of the coral holobiont is driven by the coral host, algal symbionts, and associated microbiome and is a complex trait (8). Individually, different members of populations display differing levels of stress tolerance and ability to recover after stress (39). Interestingly, we found that the adults exhibited lower overall heat tolerance, compared to their symbionts, which showed high photochemical efficiencies (0.6-0.7 at heat) across both species and populations. These patterns have been shown in other species, including adult *Porites astreoides* (40) and *Acropora palmata* (41) colonies. In these studies, adults with the same symbiont communities displayed different heat stress responses, signifying the role of the host in determining phenotypes compared to their symbionts. Drury et al. (42) also showed extensive variation in bleaching of *Acropora cerviconis* across distinct colonies with a single dominant symbiont community. Taken together, this shows that the relative contribution of the host coral or symbiont community in driving the heat tolerance response is highly variable, complex and deserves further study.

Finally, we observed relative low levels of bleaching, but advanced and rapid necrosis followed by mortality in adult corals in both populations. This is likely due to the extreme heat stress (∼ +3 ^°^C for OS and +4.5 ^°^C for PP above reported MMM) corals were exposed to. Importantly, the experimental stress coincided with corals’ summer temperature maxima (due to their collection in March), suggesting these corals may have already been exposed to peak annual temperatures. Similar patterns of rapid tissue loss under acute heat stress (+ 7 ^°^C above summer maxima) have been observed in other experimental studies (43,44) and rapid death without bleaching has also been observed in the wild and has been linked to extreme warming conditions (45). By 2100 on Ningaloo, mean temperatures are projected to rise by +1.24 ^°^C, which will exceed present summer maxima for many local reefs (4). Our results highlight the increasing vulnerability of Ningaloo corals during their critical reproductive window in the warmest summer months.

## Conclusions

Here we show that after only one generation of breeding along a relatively small geographic distance and temperature differential, selective breeding can enhance offspring heat tolerance in two widespread heat-sensitive species. However, further research is needed to identify the specific drivers of heat tolerance in adult corals and to identify which genetic markers contribute to heat tolerance, and if they are being transferred to offspring. Given future warming scenarios for Ningaloo and projections of ecological functional loss (4), we suggest that selective breeding combined with Assisted Gene Flow could be a feasible conservation tool for enhancing coral heat tolerance along Ningaloo World Heritage reef.

## Materials and Methods

### Site selection and coral collection

Three sites were initially selected along the Ningaloo Coast World Heritage Marine Park in Western Australia for coral collections. These were Oyster Stacks (OS, 22^°^ 07.869’ S 113^°^ 52.604’ E) in the northern region and Pelican Point Lagoon and Pelican Point South (PP Lagoon, 23^°^ 19.505’ S 113^°^ 46.730’ E; PP South, 23^°^ 19.416’ S 113^°^ 46.909’ E) in the southern region (Figure 1a). Colonies from the two sites in the southern region were taken as a single population.

Gravid colonies of two species (*Acropora tenuis* and *Acropora millepora*) were collected from the reef flats at a mean depth of 1.3 ± 0.5 m during high tide within a week of the anticipated onset of the autumn spawning period. For this region, spawning typically occurs within 1-10 days after full moon (07 March 2023) (46). Individual colonies were assumed to be unique genotypes by sampling individuals at least 10 m apart. The reproductive condition of each colony was assessed through visual examinations for the presence or absence of pigmented oocytes in a single colony branch sampled below the expected sterile zones following established methods (47). Fragments of ∼20 cm in width (= 20-25% of the total colony) were carefully dislodged using chisels and hammer. Ten colonies of each species were collected from Oyster Stacks, 10 from Pelican Point Lagoon, and nine from Pelican Point South (Table S1).

Coral fragments were transported in 110 L insulated coolers filled with seawater collected at the reef site, measured to be ∼27 ^°^C. Coolers were aerated individually by two battery-powered aquarium air pumps with a 150 L/hr flow rate. These were transported back to the Minderoo Exmouth Research Laboratory (MERL) either by vehicle or vessel, depending on site of collection. Colony fragments from OS and PP Lagoon were placed in two 1200 L fiberglass tanks, while those from PP South were housed in four 100 L fiberglass tanks. All tanks were supplied with 20 μm filtered seawater (FSW) at a pH of 7.7, 35 psu salinity. Tank temperatures were set at 27.1 ± 0.5 ^°^C, which was equivalent to the maximum monthly mean (March) averaged between the reef sites. A 11h:13h dark:light cycle was set, with light intensities of 50, 100, and 20 μmol photons m^-2^ s^-1^ for the OS, PP Lagoon, and PP South (hereafter referred to both as PP) tanks respectively. Light was provided by four units of the automated full spectrum dimmable 130 W reef aquarium LED lighting Orphek Atlantik iCon (Orphek, Canada) and full spectrum 72 W aquarium LED lighting Aqua Air 600 (MicMol, China) for the 1200 L and 100 L tanks, respectively. A lower light intensity was used for two sites to mitigate potential collection-induced stress. Light levels were monitored twice daily using an MQ-210 Spherical Underwater Quantum Flux sensor (Apogee Instrument, United States), while temperature was tracked continuously through the custom-built MERL control system for nine days before spawning.

### Selective breeding and larval rearing

At sunset (18:45 h), within the window of predicted spawning nights, colonies were isolated in individual 60 L polyethylene bins approximately one hour before the predicted start time for spawning (46). The bins were filled with FSW at 27.1 ^°^C, without water flow or aeration. Colonies were isolated to ensure the unintentional cross breeding did not occur in the main holding tanks. Imminent signs of spawning were monitored in all colonies (46). Released *A. tenuis* gamete bundles were collected between 19:00 h and 20:00 h from the 12^th^ to 14^th^ of March from four OS and five PP colonies. For *A. millepora*, gametes were collected between 20:30 h and 21:30 h on the 16^th^ of March from one OS and five PP colonies. Released gamete bundles were immediately skimmed from the water surface in the bins using 350 mL cylindrical polyethylene containers. Eggs and sperm from each colony were separated by gentle agitation through a 120 μm nylon mesh sieve, and eggs were washed with flow-through FSW for five to 15 minutes to completely remove remaining sperm attached to the outside of eggs. For each colony, cleaned eggs were collected into individual 2 L polyethylene bowls and concentrated sperm into individual 400 mL graduated transparent polyethylene containers. After washing, eggs and sperm were immediately moved to a controlled-environment room set to 27 ^°^C. Care was taken through this whole process to avoid cross-contamination between individuals by appropriately labeling containers with the colony identities and by rinsing any containers or materials used with a 1% diluted bleach and FSW solution.

Reproductive crosses followed such that eggs and sperm were mixed in specific combinations to create distinct families (Table S2). Families were produced by adding an equal concentration of sperm from one parental colony to an equal number of clean and isolated eggs from another colony. The density of sperm was estimated based on previous serial sperm dilutions and diluted to optimal concentrations for these species (∼10^6^) (47). A total of 44 distinct families were produced across the two species, including 22 intrapopulation crosses (OS×OS, PP×PP) and 22 interpopulation crosses (OS×PP, PP×OS), with the maternal colony identity listed first followed by the paternal colony. For *A. millepora*, the intrapopulation cross OS×OS was not produced due to insufficient gametes from unique OS colonies.

Eggs and sperm were allowed to fertilize for 3 h, with fertilization success verified by observing initial embryo cleavage at 1.5 h post-fertilization (pf) (48). When the fertilized eggs from *A. tenuis* crosses reached the four-cell stage, they were moved to separate 15 L transparent cone-shaped polycarbonate tanks (Pentair Aquatic Eco-Systems, United States) and kept at densities of ∼1-1.2 larvae/mL (15), supplied with 20 μm FSW at 27 ^°^C without photosynthetic lights given coral larvae from these species are aposymbiotic. Aeration was turned off until early gastrula stage development (> 24 h) (48). *A. millepora* cultures were reared in 800 mL clamshell-shaped polyethylene containers without flow-through. Complete daily water exchanges using 20 μm FSW at 35 psu salinity were performed on days 1 and 2 pf as per Marhaver et al. (49). *A. tenuis* larvae were motile on days 3 and 4 pf, while *A. millepora* larvae were motile on day 2 pf, consistent with the larvae development (48).

### Larval heat stress experiment

Twenty larvae per family were pipetted into 24 mm netwell inserts with a 74 μm mesh polyester membrane (Corning, United States) (Table S3). This included n=3 netwell replicates per family per temperature placed into a 6-well floating high-density polyethylene plate as described in Weeryianun et al. (28) and Quigley and van Oppen (11). These netwells were then placed into two 1200 L fiberglass tanks filled with 20 μm FSW, maintained at a pH of 7.7, 35.5 psu salinity, and under ambient room lighting. Control and heat treatments were set to 27.1 ± 0.5 ^°^C and 35.5 ± 0.5 ^°^C, respectively. At the start of the experiment, the heat treatment tank was ramped from 27.1 ^°^C using an automatic program at a rate of 0.5 ^°^C increments per hour until 35.5 ^°^C. The heat treatment temperature was chosen for comparison with previous selective breeding studies on the GBR (11, 13, 27, 28). In-tank temperatures were recorded every minute using temperature data logger HOBO 4-channel analog (Onset, United States) (Figures S1b and S1c). The number of live larvae was counted twice daily until ∼50% mortality was reached across all families in the heat treatment (19-25 March) (Figure 1b).

### Adult heat stress experiment

Multiple colonies for *A. tenuis* (OS = 10, PP = 19) and *A. millepora* (OS = 10, PP = 10) were sectioned into fragments of 5.0 ± 0.7 cm (M ± SE) in length for the adult heat stress experiment after spawning (Table S1). Colonies were fragmented into small nubbins using a Gryphon Aquasaw diamond band saw (Gryphon Corporation, United States). At the start of the experiment, all colonies showed a healthy appearance (i.e., minimal bleaching) and care was taken to cut similarly sized nubbins per colony. Individual fragments were immediately glued onto aragonite plugs (19 mm crown and 11 mm base, Ocean Wonders, United States) using cyanoacrylate adhesive (Gorilla Glue Company, United States). Fragments were labeled with their corresponding colony identification onto each nubbin and placed onto custom-built polyethylene plug holders. For acclimation, plug holders were placed into 50 L acrylic tanks supplied with 20 μm FSW, maintained at 27.1 ± 0.5 ^°^C, pH of 7.7, and 35 psu salinity. Each tank was equipped with a wavemaker pump and an air stone for aeration and water mixing, and the light cycle was set to an 11h:13h dark:light regime with a gradual 6 h increase to a maximum PAR of 60 μmol photons m^-2^ s^-1^ provided by one unit of the automated full spectrum dimmable130 W reef aquarium LED lighting Orphek Atlantik iCon (Orphek, Canada).

After 8-9 days of acclimation, fragments were rearranged into a randomized design across eight tanks such that each colony genotype was represented with n = 3-4 fragment replicates per colony per temperature across replicate tanks for control and heat. Control and heat treatments were set at 27.1 ± 0.5 ^°^C and 31.0 ± 0.5 ^°^C, respectively. In the heat treatment, 27.1 ^°^C water was automatically ramped up in 0.5 ^°^C increments per hour until 31.0 ^°^C, matching the maximum sea surface temperature that corresponded to the DHW values during the 2011 MHW in Western Australia (20). Treatments were maintained until ∼50% mortality was averaged across all genotypes per species in the heat treatment. In-tank temperatures were recorded every minute using the temperature data logger HOBO 4-channel analog (Onset, United States) (Figures S1a and S1b), and fragments were fed daily after dusk with one scoop of Vitalis Mixed Reef Food (Vitalis Aquatic Nutrition, United Kingdom).

Coral fragments were photographed daily in separate tanks with the same treatment temperatures. Photographs were used to determine survival, tissue necrosis, and bleaching scores. Photos were taken using an Olympus Tough TG-6 digital camera (Olympus, Japan) positioned 50 cm from the tanks with fixed settings (ISO 200, focal lens of 25 mm, focal ratio of *f*2.8, and shutter speed at 1/50 s) and tank illumination (one unit of 72 W aquarium LED lighting Aqua Air 600 set at 50% white). For survival determination, live fragments were scored “1” and dead fragments were scored “0”. Dead was defined as bare skeleton with or without microalgae overgrowth. Percent necrosis was measured using the surface area tool in the Fiji software (50) as described in Quigley et al. (27). Bleaching was determined by assigning scores to fragments relative to the brown hue (D1-D6) from the Coral Health Chart used as proxy for changes in symbiont density and chlorophyll-a content (CoralWatch, Australia; 51). Bleaching scores of D1 (white) are indicative of a bleached fragment and D6 (brown) of a non-bleached healthy fragment. Scores were assigned to three random points along a vertical axis of each fragment and averaged. Photochemical efficiency of photosystem II in a light-adapted state was measured by the effective quantum yield (ΔF/Fm’) (44) at the start of peak light intensity (10:00 h) using a DIVING-PAM fluorometer (Walz, Germany) using the following settings: measuring light intensity = 3, saturation pulse intensity = 8, saturation pulse width = 0.8 s, gain = 6 and damping = 2. Measurements were taken consistently 10 mm from the coral tissue and ∼20 mm above the fragment base. Physiological responses were measured until an average of 50% species mortality was reached during the period from 25-31 March (Figure 1c).

### Temperature metrics

To determine thermal regime differences at OS and PP and use as proxy for coral heat tolerance in selective breeding, sea surface temperature (SST) metrics were computed for two timeframes:

1. before recent MHW events (01 January 1985 to 30 September 2010, “historical”) and 2) during and after MHWs (01 October 2010 to 29 December 2022, “post-MHW”) (2).

Metrics based on predictors for coral bleaching resistance and heritability of heat tolerance (see 11) included: mean annual SST (SST_mean), standard deviation of annual SST (SST_stdev), diurnal SST range (DTR), frequency of thermal anomalies (SSTA_Freq), and cumulative thermal stress in degree heating weeks (TSA_DHW) (Table S4). Variation of SSTA_Freq and TSA_DHW were not included due to insufficient data points for statistical analysis.

Metrics were derived from NOAA’s CoralTemp SST product (v 3.1) (52), Coral Reef Temperature Anomaly Database (CoRTAD, v 6) (53), and IMOS’s Himawari-8 L3C (54), which offer the highest spatial resolution for Western Australia. Data for the bounded area of interest (i.e., latitudes 23^°^ 33.924’ S to 21^°^ 46.923’ S and longitudes 113^°^ 29.082’ E to 114^°^ 0.078’ E) were downloaded in January 2023 as netCDF files via the THREDDS data server using Python (v 3.9.14 64-bit).

## Statistical analyses

Temperature metrics were calculated in R (v 4.2.2; 55). For monthly SST timeseries, we averaged monthly SST from daily SST. For SST_mean, we averaged daily SST for each year at each location to obtain the mean annual SST across all years. Similarly, SST_stdev was calculated by averaging the standard deviation of annual SSTs. DTR was obtained by subtracting the minimum daily SST from the maximum daily SST and averaging across available years (2016 to 2022). SSTA_Freq and TSA_DHW were derived directly from the CoRTAD dataset (53). Except for DTR, all metrics were calculated for the historical and post-MHW periods.

Data normality and homogeneity were checked using Shapiro-Wilk and Levene’s tests, with ‘Shapiro.test’, ‘leveneTest’, and ‘qqnorm’ from the R package ‘stats’ (v 4.2.2) and ‘car’ (v 3.1-2; 56). Statistical differences were tested using the non-parametric Wilcoxon’s Rank Sum Test (‘wilcox.test’ from the R package ‘stats’), with a significance level set at 0.05. Temperature metrics were visualized using ‘ggplot2’ (v 3.4.4; 57).

To assess survival differences in adult fragments between two populations per species, Kaplan-Meier (KM) survival curves were used. Survival data was transformed into individual mortality events for KM models. Survival probabilities for control and heat treatments were determined at seven time points. Pairwise comparisons of survival curves were performed using a post-hoc Log-Rank test with P values corrected using the Bonferroni method (58). Survival probability with the interaction factor (INT) (temperature*population) was used for comparisons. Survival curves were plotted for each reef and species in the control and heat treatments. R packages ‘survival’ and ‘survminer’ were used for this analysis (v 3.5-7; 59; v 0.4.9; 60). The same procedure was applied to evaluate larval survival differences by cross type and parental origin.

Differences in percent necrosis, bleaching scores, and effective quantum yields at the experimental endpoint were evaluated against temperature treatment and population using generalized linear mixed models (glmms) from the R packages ‘lme4’ (v 1.1-35.1; 61), ‘glmmTMB’ (v 1.1.8; 62), and ‘MASS’ (v 7.3-60.0.1; 63). Temperature and population were converted into an interaction factor (INT), set as a fixed factor in all models. Genotype was the only significant random effect after testing multiple factors. Zero and one inflation were non-significant and dropped from the models. Model assumptions were checked using diagnostic plots and tests from ‘stats’ (v 4.2.2; 57) and ‘DHARMa’ (v 0.4.6; 64). Percent necrosis was converted into proportion in a betaGLMM. A penalized quasi-likelihood (Gaussian distribution) glmm (PQLglmm) was used for effective quantum yields and a glmm with negative binomial distribution for bleaching scores. Post-hoc Tukey’s pairwise comparisons between temperature and population were run on models’ outputs using the function ‘ghlt’ from the ‘multcomp’ package (v 1.4-25; 65) with P values corrected using the Bonferroni method.

Data for each adult and larval response was summarized with ‘dplyr’ (v 1.1.4; 66) and plotted using ‘ggplot2’ (v 3.4.4; 57). Survival of individual larval families was plotted using ‘ggplot2’, and families with significant declines in survival in the heat treatment relative to control were identified using a parametric one-tailed t test using the t.test function from ‘stats’ package (55). P-values were corrected for false discovery rate using ‘stats’ (v 4.2.2; 55). All statistical analyses were carried out using R (v 4.2.2; 55).

